# The Deep(er) Roots of Eukaryotes and Akaryotes

**DOI:** 10.1101/2020.01.17.907717

**Authors:** Ajith Harish, David A. Morrison

## Abstract

Locating the root-node of the “tree of life” (ToL) is one of the hardest problems in phylogenetics^1^. The root-node or the universal common ancestor (UCA) divides the descendants into organismal domains^2^. Two notable variants of the two-domains ToL (2D-ToL) have gained support recently^3,4^, though, Williams and colleagues (W&C)^4^ claim that one is better supported than the other. Here, we argue that important aspects of estimating evolutionary relatedness and assessing phylogenetic signal in empirical data were overlooked^4^. We focus on phylogenetic character reconstructions necessary to describe the UCA or its closest descendants in the absence of reliable fossils. It is well-known that different character-types present different perspectives on evolutionary history that relate to different phylogenetic depths^5–7^. Which of the 2D-ToL^2,4^ hypotheses is better supported depends on which kind of molecular features – protein-domains or their component amino-acids – are better for resolving the common ancestors (CA) at the roots of clades. In practice, this involves reconstructing character compositions of the ancestral nodes all the way back to the UCA^2,3^.

## 1. Introduction

Models of character evolution are essential to determine the evolutionary relationships of organisms. Phylogenetic models that employ protein structural-domains as characters place Asgards as sister to other archaea, and archaea sister to bacteria in the “tree of life” (ToL)^1–3^. Whereas several analyses that employ amino-acids as characters fail to resolve the archaeal radiation or to identify a distinct ancestor of archaea^4^. In a recent study, Williams and colleagues (W&C)^4^ compared the performance of several character-evolution models to evaluate which one of the ToL hypotheses is better supported. The authors tested the performance of several substitution models for amino-acid characters using empirical data, but models for protein-domain characters with simulated data.

W&C rely on (i) simulated data to reject a robust phylogeny inferred from empirical data (Fig.1a) that supports the evolutionary kinship of eukaryotes and akaryotes (*akaryote 2D-ToL*)^1–3^; and (ii) the so-called bacterial rooting to interpret a partially resolved, unrooted-ToL (Fig.1b), asserting that Asgard archaea are the closest relatives of eukaryotes (*eocyte 2D-ToL*)^4^. Both are questionable since (i) simulated data neither reproduce nor represent empirical distributions; and (ii) unresolved trees obscure evolutionary relationships. In this article, we argue that (i) W&C have overlooked important aspects of assessing phylogenetic signal in empirical data, and (ii) it may be premature to reject a well-supported phylogeny based on simulated data.

**Fig. 1.**
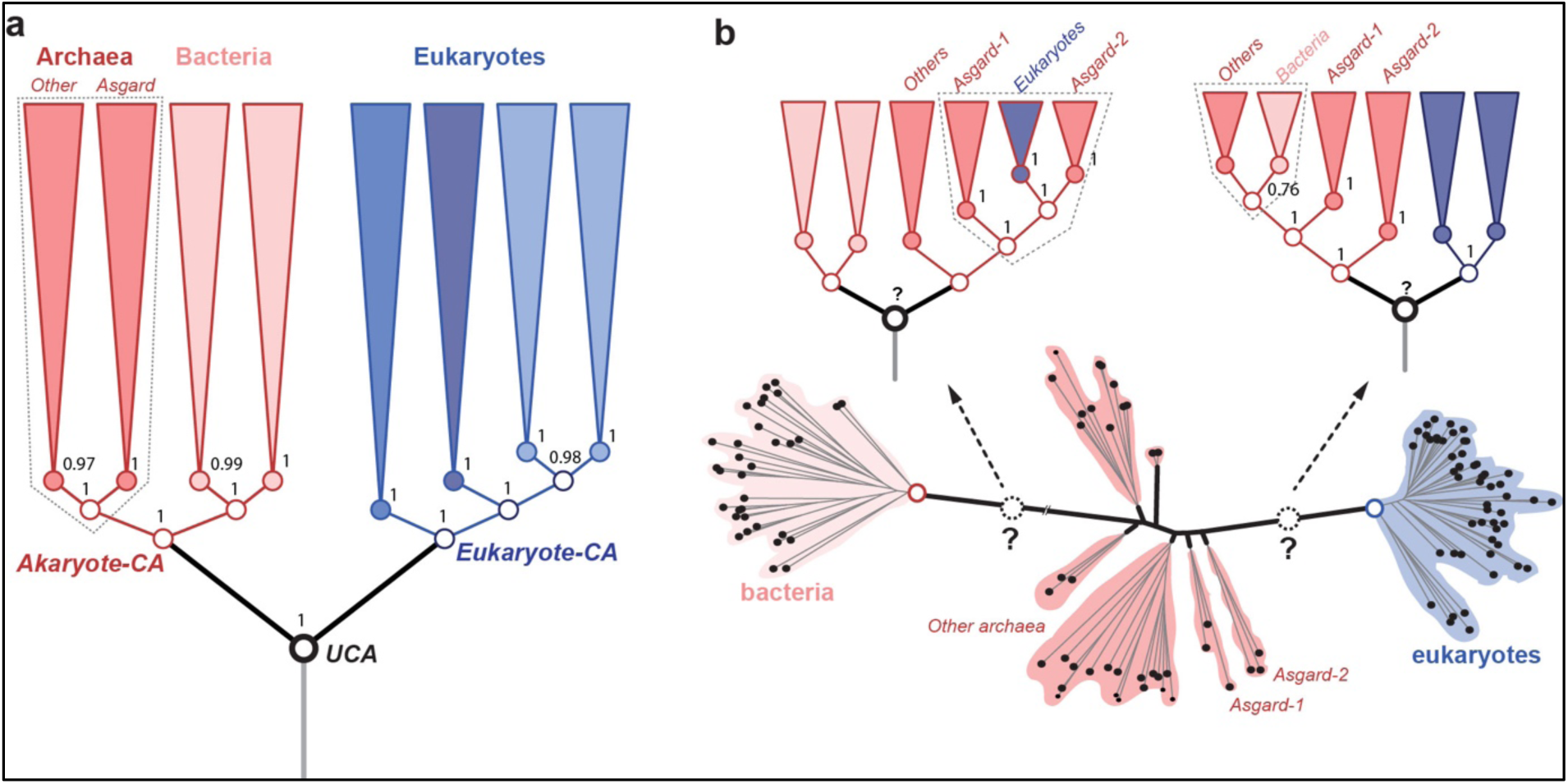
Different 2D “tree of life” (2D-ToL) variants supported by different types of molecular characters using the best-fitting probability models^1,4^. **a**, The rooted tree (phylogeny) inferred by estimating the evolution of species-specific changes in protein-domain composition. Directional character-evolution models place the root between eukaryotes and akaryotes. Named groups of organisms, including Asgardarchaeota are resolved into clades (i.e. a single ancestor). The Asgard archaea are sister to all other archaea, with euryarchaea being the closest relatives. The phylogeny shown is a condensed form obtained after collapsing the clades of the full tree in ref^1^. **b**, The unrooted tree inferred by estimating the evolution of protein/gene-specific changes in amino-acid composition. The unrooted-tree is the same as in fig. S8d in W&C^4^. The group archaea, and Asgard archaea are unresolved; and a distinct archaeal ancestor is absent. Time-reversible character evolution models cannot identify the root (UCA) as well. Alternative rootings polarize the branching order in opposite directions implying incompatible relationships among the major organismal clades. Regardless of the rooting, neither Asgard archaea nor archaea as a whole can be resolved as a monophyletic group. Further, Argards do not share a unique common ancestor with other archaea. Even the best-fitting amino-acid evolution models cannot resolve the archaeal radiation despite employing thousands of genes^4^. The poor resolution of archaea is seen in virtually all trees, with or without inclusion of long-branches of bacteria. In such ambiguous cases, “character polarization” as in (**a**) is likely to be efficient, rather than the more commonly used “graphical polarization” of unrooted trees. Clade support is indicated for key groups as (**a**) Bayesian posterior probability, (**b**) bootstrap percentage.

## 2. Results and Discussion

### 2.1 Which molecular feature is a better phylogenetic character?

Reversibility of amino-acid replacements due to biochemical redundancy makes determining character compositions of ancestral nodes ambiguous, as character polarity is ambiguous. This has been a sticking point for locating a distinct archaeal-CA to resolve the archaeal radiation. This is routinely seen as a conspicuous absence of the archaeal-CA as well as UCA in unrooted trees (e.g. Fig. 1b), inferred using time-reversible models of character evolution^4,8,9^. Without a distinct node to unite the archaeal branches, the archaea are unresolved, whereas eukaryotes and bacteria are resolved so that their CA-nodes are discernable.

Protein structural-domains are biochemically non-redundant, unlike amino-acids, and have proven to be excellent “genomic characters” ^1,2^ that support a robust *akaryote 2D*-*ToL* (Fig 1a). Though undervalued, they afford many conceptual and technical advantages over amino-acids for reliable phylogenetic modeling^1,7,10^ and estimating ancestral compositions^2,3,11^:

- Substitutions between structural-domains do not occur, unlike amino-acid replacements, since each domain defines a distinctive biochemical function^1^ (Fig. 2a).
- The natural bias in gain/loss rates, arising from the difficulty of parallel gains and the relative ease of parallel losses, is useful for implementing directional (rooted) character-evolution models^3,11,12^.

A key advantage of non-redundant characters is that estimating ancestral compositions and evolutionary paths of individual characters is much less ambiguous. In addition to identifying the root-nodes, an added benefit of the built-in directionality is that mutually exclusive evolutionary fates of individual features – inheritance, loss or transfer – can be resolved efficiently using directional-evolution models^1,12,13^.

**Fig. 2.**
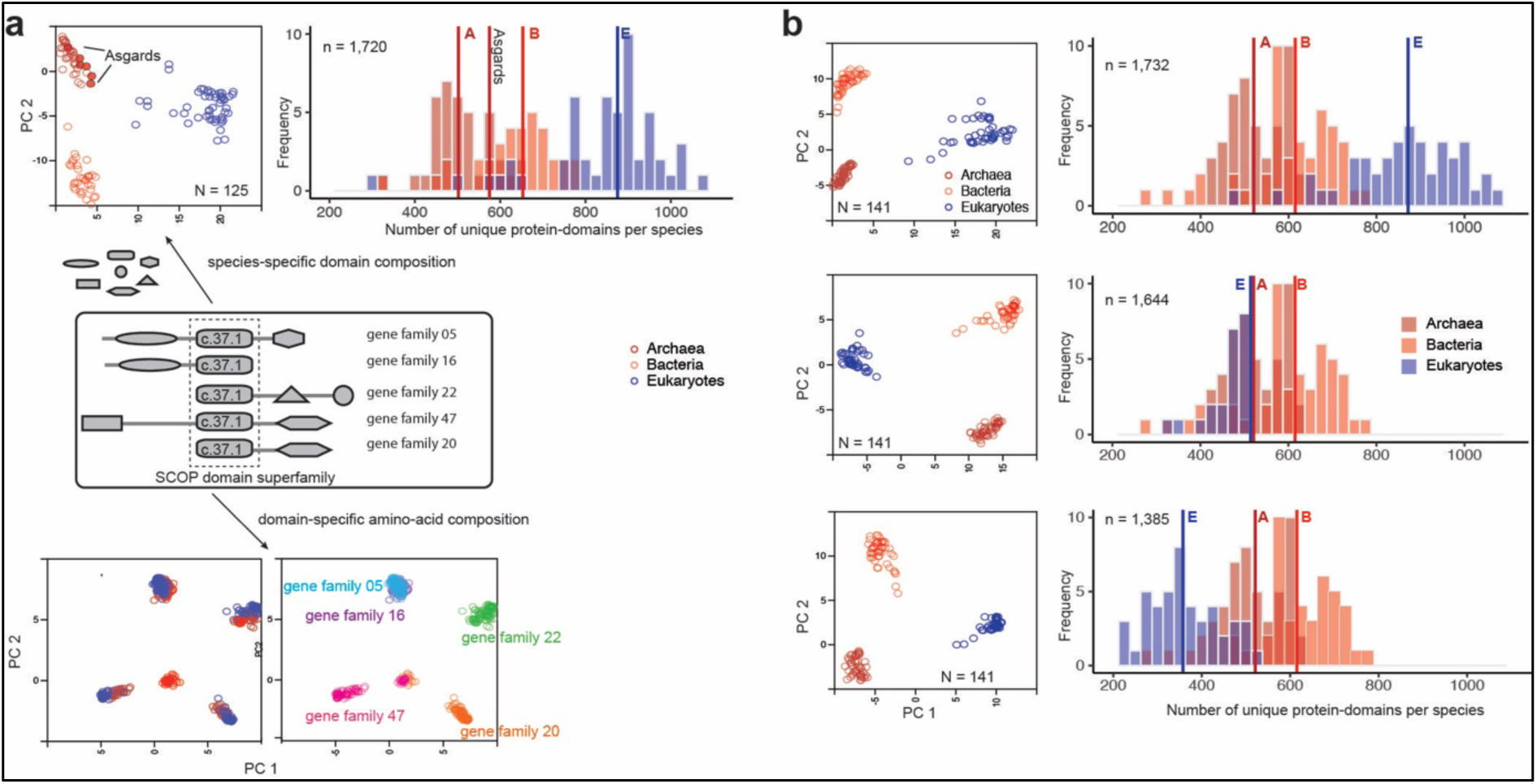
Compositions of unique protein-domains identify with organismal families whereas amino-acid compositions of individual domains relate to gene families. **a**, Protein-domains are considered to be independent evolutionary units with a distinct tertiary fold, amino-acid sequence and biochemical function. The Majority of proteins are multi-domain proteins formed by duplication and recombination of domain units. Covariation of protein-domain composition among the 125 species sampled in C&W^4^ (top) was compared by principal components analysis (PCA). Each circle in the PCA projection (top left) is a distinct species, defined by a species-specific domain cohort. Asgards are highlighted as filled circles. The frequency distribution (top right) shows the number of distinct protein-domains per species. Vertical intersecting lines in the histograms are the median numbers of protein-domains. Protein-domain composition is characteristic of clades of species (top left). In contrast, covariation of amino-acid composition (bottom) in a single-domain (super)family is not clade-specific, but gene family-specific. Sequences alignments of a single-domain (c.37.1) shared by 5/50 concatenated orthologous gene-families from 125 species were sampled for the PCA projection. **b**, Effects of severe perturbation of the domain composition in recovering clade-specific distributions was tested in a sample of 141 species. Although it is common to suspect that the rooting between akaryotes and eukaryotes could be biased due to a larger domain cohort in eukaryotes^4^, it is not the case^2,3,11^. Diversity of clade-specific domain composition (top right) measured simply as the number of protein-domains^4^ is a poor descriptor of heterogeneity, and can be misleading. Clades are grouped by covarying “protein-domain types”, but not by numbers alone. The rooting is stable and the tree topology is virtually identical even after reducing the eukaryote cohort by 1/3rds (middle) or 2/3rds (bottom)^13^ of the original composition^2^. Description of the PCA projections and frequencies are the same as in (**a**).

To be clear, unrooted trees are not phylogenies *per se* since the absence of root-ancestor(s) obscures ancestor-descendant polarity and *phylogenetic relatedness*^14,15^. Since identifying the closest relatives of extant groups is the same as determining the closeness of their common ancestors, time-reversible models and unrooted trees remain ineffective tools. Thus, regardless of the gene-aggregation and tree-reconciliation approach used for estimating a consensus unrooted tree^4^, the location of the archael-CA or UCA remains ambiguous (Fig. 1b). Support from fossils or other sources are not reliable, despite claims to the contrary^4^. Likewise, predicting the origins of single-domains or single-genes by estimating amino-acid (or nucleotide) compositions also remains ambiguous (reviewed in refs^1,6,7^). A sobering revelation is that some datasets/models may be of little use or relevance to resolve questions of deep time evolution — this is sad but true.

### 2.2 Will more complex models minimize uncertainties?

W&C^4^ argue that (i) directional-evolution models^11,12^ may be unsuitable to predict the unique origin of homologous protein-domains; and (ii) the *akaryote 2D-ToL*^*2,3*^ is an unsatisfactory explanation of the clade-specific compositions of protein-domains (Fig. 2). Their arguments seem to imply that phylogenetic signal can be recovered only by modeling evolution of amino-acid composition. However, the fact that even the best-fitting substitution models are inadequate^4^, despite ever increasing model complexity, suggests that different protein-domain families may require different, but incompatible substitution models (Fig. 2a).

The KVR^12^ model is an extension of the Mk [Markov *k* states] model^16^, a generic probability model for discrete-state characters. A variant at *k* >=20 is suitable for modeling evolution of amino-acids or copy numbers of gene or protein-domain families. While time-reversible variants produce unrooted trees, such directional models consistently recover a 2D phylogeny (Fig. 1a) in which akaryotes are the closest relatives of eukaryotes^1,2,13^. The KVR model assumes that the root-ancestor has a different character composition than the rest of the tree, which is essentially an irreversible acyclic process. This is fully consistent with the idea that, on a grand scale, the “tree of life” describes broad generalizations of singular events and major transitions underlying striking sister clade differences. Since parallel evolution of homologous protein-domains or distinct domain permutations is very rare, the KVR model adequately captures the evolution of unique features.

This assumption is also consistent with the idea that the idiosyncratic compositions of homologous protein-domains (Fig. 2) is a characteristic of the clades^1–3^. In contrast, amino-acid compositions in single-domain families are not (Fig. 2a). The systematic covariation of homologous domains among the clades is best explained as phylogenetic effect. Consequently, the akaryote 2D-ToL (Fig.1a) was consistently recovered with robust support for the major clades regardless of the taxonomic/protein-domain diversity sampled (Fig. 2b), and regardless of the model complexity^1–3,11,13^.

The KVR model is an optimal explanation of the evolution of clade-specific composition of homologous features. Complex variants of the KVR model that account for rate variation among both characters and branches also consistently recovered the akaryote 2D-ToL despite significantly different model-fits^1^. More complex models are available, such as the no-common-mechanism model^17^, an extremely parameter-rich model that allows each character to have its own rate, branch length and topology parameters. Even more complex models can be implemented, which assume that the tempo and mode of evolution changes at each internal node along the phylogeny^4^. However, such over-specified models may not be optimal for generalizing the evolutionary process; and may over-fit observed patterns - a form of model misspecification. That said, it remains to be seen whether more complex models perform better with empirical datasets.

## 3. Data and Methods

### 3.1 Data sources

Proteome sequences (predicted protein cohorts from genome sequences) were obtained from recently published studies^4,13^. Homologous protein structural-domains were identified using the homology assignment tools provided by the SUPERFAMILY database as in previous studies. Briefly, each proteome was queried against the hidden Markov model (HMM) library of homologous protein-domains defined at the Superfamily level in the SCOP (Structural Classification of Proteins) hierarchy. The taxonomic diversity of sequenced genomes and the number of unique protein-domains identified for each species is as follows:

| Number of species sampled per clade | Number of unique protein-domains |
| --- | --- |
| 125 (Archaea: 39; Bacteria: 33; Eukarya: 52)<sup>4</sup> | 1,720 |
| 141 (Archaea: 47; Bacteria: 47; Eukarya: 47)<sup>13</sup> | 1,732 |

### 3.2 Statistical analysis

Descriptive statistics of protein-domain compositions for each taxonomic sampling including the frequency distribution and median number of protein-domains for each clade (Archaea, Bacteria and Eukarya) were estimated and visualized using the ggplot2 package in R 3.6.2. Covariation of clade-specific protein-domain composition as well as domain-specific amino-acid composition was compared using principal component analysis (PCA). Components were generated by an eigenvector decomposition of the character matrix. PC scores were based on percentage identity of character compositions.

## Acknowledgements

We thank Tom Williams for kindly providing the proteome sequences used in their study and for answering our questions.

## Author contributions

A.H. designed the study and analyzed the data. A. H. and D.A.M wrote the paper.

## Competing interests

The authors declare no competing interests.

